# Assessing neurophysiological changes associated with combined transcranial direct current stimulation and cognitive emotional training for treatment-resistant depression

**DOI:** 10.1101/688317

**Authors:** Stevan Nikolin, Donel Martin, Colleen K. Loo, Brian M. Iacoviello, Tjeerd W. Boonstra

## Abstract

Transcranial direct current stimulation (tDCS), a form of non-invasive brain stimulation, is a promising treatment for depression. Recent research suggests that tDCS efficacy can be augmented using concurrent cognitive emotional training (CET). However, the neurophysiological changes associated with this combined intervention remain to be elucidated. We therefore examined the effects of tDCS combined with CET using electroencephalography (EEG). A total of 20 participants with treatment resistant depression took part in this open-label study and received 18 sessions over 6 weeks of tDCS and concurrent CET. Resting-state and task-related EEG during a 3-back working memory task were aquired at baseline and immediately following the treatment course. Results showed an improvement in mood and working memory accuracy, but not response time, following the intervention. We did not find significant effects of the intervention on resting-state power spectral density (frontal theta and alpha asymmetry), time-frequency power (alpha event-related desynchronization and theta event-related synchronisation), or event-related potentials (P2 and P3 components). We therefore identified little evidence of neurophysiological changes associated with treatment using tDCS and concurrent CET, despite significant improvements in mood and near transfer effects of cognitive training to working memory accuracy. Further research incorporating a sham controlled group may be necessary to identify the neurophysiological effects of the intervention.

## INTRODUCTION

Major depressive disorder (MDD) is a debilitating mental illness with a lifetime prevalence of 12-20% (Mrazek *et al*., 2014). Transcranial direct current stimulation (tDCS), a mild, non-invasive method of brain stimulation, has shown promising therapeutic potential for the treatment of depression (Loo *et al*., 2012; Brunoni *et al*., 2013; Brunoni *et al*., 2016; Brunoni *et al*., 2017). However, results from a recent large controlled trial suggested that such antidepressant effects of tDCS when given alone are modest (Brunoni *et al*., 2017), raising the need for research into improving tDCS clinical outcomes. Combining tDCS with concurrent cognitive training (e.g., cognitive emotional training, designed to activate brain regions associated with cognitive control and emotion processing) appears to be a promising method to further boost treatment outcomes (Martin *et al*., 2018). The neural pathways by which this novel combined intervention improves mood remains unknown. An examination of the neurophysiological changes associated with this approach may shed light on its mechanisms of action and pave the way for further augmentation of the therapy.

tDCS for depression involves the delivery of an electrical current, typically between 1-2.5 mA, using a positively charged electrode placed on the scalp over the left dorsolateral prefrontal cortex (DLPFC) and a negatively charged electrode on a contralateral frontal region (Loo *et al*., 2018). Though research suggests that tDCS is an effective treatment in MDD in general (Mutz *et al*., 2019), the results in treatment resistant depression appear less promising (Blumberger *et al*., 2012; Palm *et al*., 2012). Indeed, a meta-analysis of individual patient data found treatment resistance to be associated with reduced tDCS efficacy (Brunoni *et al*., 2016).

To improve therapeutic efficacy in people with treatment resistant depression, some trials have combined tDCS with cognitive training (Brunoni *et al*., 2014; Segrave *et al*., 2014). It has been suggested that tDCS interacts with ongoing neuronal activity to boost synaptic plasticity within activated regions (Kronberg *et al*., 2017; Kronberg *et al*., 2019). The administration of behavioural tasks during tDCS may thus be used to pre-activate relevant cortical regions and augment the neuromodulatory potential of tDCS by improving the functional specificity of stimulation (Bikson & Rahman, 2013). Cognitive emotional training (CET) is a form of computerised cognitive training which was designed to activate cognition- and emotion processing-circuits implicated in MDD to treat depressive symptoms (Iacoviello *et al*., 2014; Iacoviello & Charney, 2015). Specifically, it involves training using an emotional working memory paradigm that aims to simultaneously activate brain regions in the cognitive control (i.e., prefrontal cortex/DLPFC)) and affective networks (i.e., amygdala) to improve emotion regulation. A recent open-label pilot study conducted by our group combined tDCS with concurrent CET, and reported a 41% response rate in treatment resistant participants (Martin *et al*., 2018). The neurophysiological changes associated with this intervention, however, remain to be elucidated. Determination of such changes may provide insight into the mechanisms underlying the antidepressant response and inter-individual predictors which can assist with the further development and targeting of treatment.

Electroencephalography (EEG) allows for non-invasive assessment of brain activity and has been used to gain further insights into the functional changes associated with antidepressant response (Iosifescu, 2011). Given the paucity of research on the cumulative effects of tDCS on EEG outcomes, we investigate several EEG measures that have shown relevance in the broader depression and working memory literature. EEG markers acquired at rest can provide valuable information regarding severity of depression symptomatology. Frontal alpha asymmetry, in which higher alpha power is observed in the left compared to the right prefrontal cortex (Debener *et al*., 2000; Gordon *et al*., 2010; Stewart *et al*., 2010), has been inversely correlated with depression scores (Diego *et al*., 2001). Although recent research has questioned the diagnostic value of frontal alpha asymmetry as a biomarker for depression (Van Der Vinne *et al*., 2017), alpha asymmetry has been associated with response to antidepressant medications (i.e. selective serotonin reuptake inhibitors) in females, and may therefore have some prognostic value (Arns *et al*., 2016). As the combined tDCS and CET intervention involves concurrent stimulation and activation of left prefrontal regions, it is possible this measure may additionally show changes following treatment. In addition to alpha band activity, resting-state frontal theta power may also provide a marker for depression (Iosifescu, 2011; Bailey *et al*., 2018). Several studies have reported elevated frontal theta as a positive predictive factor for antidepressant response (Pizzagalli *et al*., 2002; Spronk *et al*., 2011; Rentzsch *et al*., 2014; Arns *et al*., 2015; Koo *et al*., 2017), although another study suggested that mood changes are negatively correlated with frontal theta activity (Knott *et al*., 2000).

Task-related EEG measures, such as event-related potentials (ERPs) and time-frequency power, have also shown sensitivity to the neurophysiological effects of tDCS as compared to behavioural outcomes on a working memory task (Nikolin *et al*., 2018b). Two ERP components, P2 and P3, have been found to be particularly sensitive to the effects of tDCS during a task of working memory, with both increasing in amplitude following stimulation of the prefrontal cortex (Keeser *et al*., 2011). The P2 amplitude reflects sustained attention and initiates context updating during working memory tasks (Kemp *et al*., 2006; Luu *et al*., 2014; Yuan *et al*., 2016; Vilà-Balló *et al*., 2018), and its amplitude has been observed to be positively correlated to working memory performance (Han *et al*., 2013). The P3 component appears to reflect higher order executive processes within the frontoparietal cortical network (Polich, 2007; Brydges & Barceló, 2018), and has also been positively correlated with working memory capacity and task performance (Gevins & Smith, 2000; Dong *et al*., 2015; Gajewski & Falkenstein, 2018; Nikolin *et al*., 2018b). Event-related synchronisation (ERS) and desynchronization (ERD during working memory cognitive processes have been mainly found in the theta and alpha band (Gomarus *et al*., 2006; Missonnier *et al*., 2006; Zhao *et al*., 2017; Wianda & Ross, 2019). In healthy individuals, anodal tDCS of the prefrontal cortex has been shown to improve working memory performance and increase event-related alpha and theta power (Zaehle *et al*., 2011). Frontal theta ERS during working memory has been linked to allocation of attention to task-relevant stimuli (Missonnier *et al*., 2006). Both alpha ERD and theta ERS become more pronounced with increasing cognitive load (Krause *et al*., 2000; Stipacek *et al*., 2003; Klimesch *et al*., 2004; Klimesch *et al*., 2006), and frontal theta activity has been found to be reduced following active compared to sham tDCS during memory retrieval in depressed participants (Powell *et al*., 2014).

The clinical and cognitive outcomes of tDCS combined with concurrent CET for participants with treatment resistant depression are reported in greater detail in a previous report from our group (Martin *et al*., 2018). Here, we sought to understand possible mechanisms underlying treatment effects of this novel intervention by examining neurophysiological changes using EEG. We hypothesised that tDCS combined with CET would restore cognitive control network functioning, observable as an improvement in working memory performance and a reduction in frontal alpha asymmetry and frontal theta recorded during resting-state EEG. Additionally, we hypothesised that the intervention would increase the amplitude of event-related EEG components P2 and P3. Finally, we anticipated a reduction in alpha ERD and theta ERS during the working memory task, indicative of improved cognitive processing efficiency.

## MATERIAL AND METHODS

### Participants

We recruited twenty participants with treatment resistant depression, of which 10 were included in the sample analysed to investigate the clinical and cognitive efficacy of tDCS combined with CET (Martin *et al*., 2018). All participants were screened by a study psychiatrist and met DSM-IV criteria for a major depressive episode (APA (American Psychiatric Association), 1994). Additional inclusion criteria included: a score on the Montgomery-Asperg Depression Rating Scale (MADRS; Montgomery and Asberg 1979) ≥ 20, treatment resistance as defined by failure to respond to at least two adequate courses of antidepressant medications, and aged between 18 and 65 years old. Participants were excluded from the study if they had a neurological illness, were diagnosed with a psychotic disorder, had alcohol or illicit substance abuse, failed to respond to a course of electroconvulsive therapy in the current episode, were a high suicide risk, or regularly used benzodiazepines. Participants were not permitted to change medications or their dosages in the four weeks preceding the study and for the duration of the combined tDCS and CET intervention. Written informed consent was provided by all participants, and the study was approved by the UNSW Human Research Ethics Committee.

### Procedure

Participants received a course of 18 sessions of tDCS combined with CET, delivered three times per week for six weeks, as described in Martin *et al*. (2018). Neurophysiological outcomes were assessed pre-treatment (BASELINE) and at the completion of 18 sessions of the intervention (POST) in a within-subjects open label study design. EEG was recorded during 5 min of eyes closed resting-state activity, followed by 5 min of eyes open resting-state activity, and finally 7 min of task-related activity during a visual 3-back working memory task (Fig. 1A). During the eyes closed condition, participants were asked to close their eyes until the experimenter informed them that the EEG recording was complete. For the eyes open condition, participants were asked to focus their gaze on a fixation cross displayed on a computer screen.

**Figure 1.**
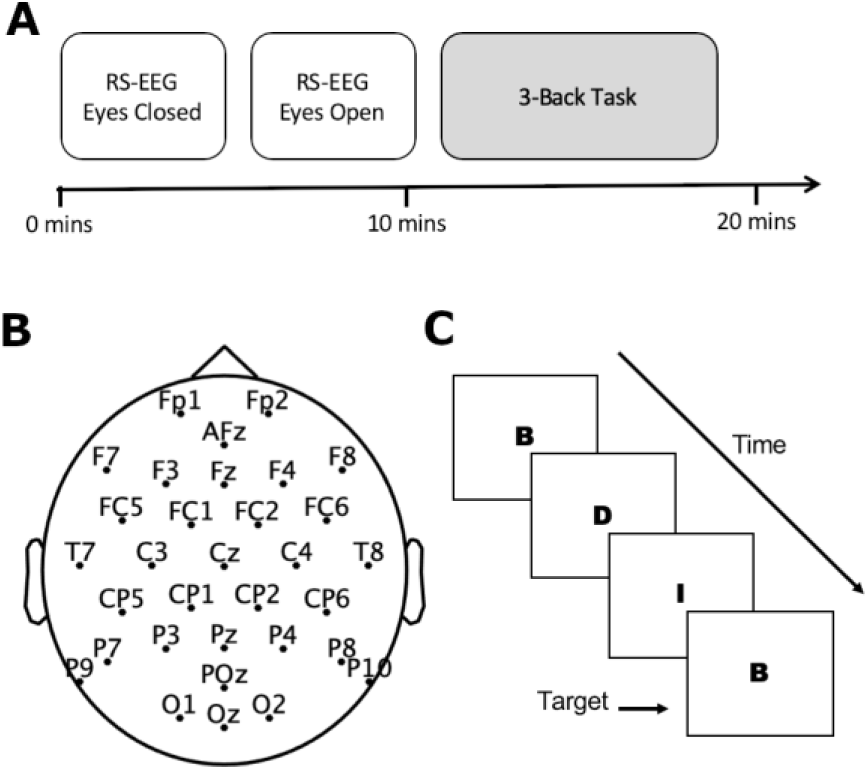
Experiment Details. **A)** Illustration of the EEG protocol and timeline. Participants completed EEG testing at baseline, and following 18 sessions of tDCS combined with CET; **B)** The EEG layout for 32 water-based electrode recording channels; **C)** For the 3-back task, participants were asked to respond when the letter currently on the screen matched the letter presented three trials previously. RS-EEG: resting-state EEG.

### Transcranial direct current stimulation

An Eldith-DC Stimulator (NeuroConn GmbH, Germany) was used to deliver a 2 mA current for 40 minutes each session. This duration was chosen to ensure that participants were stimulated for the entirety of the CET intervention, and has been demonstrated to be safe and tolerable in previous studies (Bolognini *et al*., 2011; Di Lazzaro *et al*., 2014; Gluck *et al*., 2015; Hamner *et al*., 2015). Anodal tDCS was applied using a 5 × 7 cm (35 cm^2^) rubber electrode placed on the scalp directly over the left DLPFC (F3 using the 10-20 EEG system). An extracephalic 10 × 10 cm (100 cm^2^) return cathode was placed on the participant’s right shoulder. An open-label pilot investigation found this montage resulted in more rapid mood improvement relative to a bifrontal (F3-F8) electrode placement (Martin *et al*., 2011), though its efficacy is yet to be confirmed in a randomised control trial. Computational modelling suggests that this montage increases the activation of limbic regions, such as the anterior cingulate cortex, as compared to bifrontal tDCS electrode placements commonly used for depression (Bai *et al*., 2014). Sponges soaked in saline were used improve conductivity between the electrodes and skin, and thereby minimise the risk of skin lesions (Loo *et al*., 2011).

### Cognitive emotional training

The Emotional Faces Memory Task (EFMT) was used to provide CET, and is described in greater detail in Iacoviello *et al*. (2014). In EFMT, participants observe a series of faces depicting various emotions (sadness, happiness, surprise and disgust) on a computer screen and respond (yes/no) whether the emotion observed matched the emotion presented several (a set value, “n”) trials previously, thus combining emotion processing and n-back working memory components. The task difficulty was adjusted according to participant performance by increasing or decreasing the number of emotional faces participants had to maintain in memory (n); task difficulty also progresses throughout an EFMT session as the emotional intensity observed on the faces decreases.

### Electroencephalography data acquisition

Continuous EEG data was acquired using 32 water-based EEG recording channels and a TMSi Refa amplifier (TMS International, Oldenzaal, Netherlands; see Fig. 1B). EEG processing and analysis was conducted using custom-developed MATLAB scripts (v.R2019a; MathWorks) and the Fieldtrip toolbox (Oostenveld *et al*., 2011). All scripts used for EEG processing and calculation of neurophysiological measures are available at the following link (https://github.com/snikolin/tDCSandCET).

EEG data were sampled at 1024Hz, and filtered using a bandpass filter (0.5–70 Hz) and a notch filter at 50 Hz to remove electrical line noise. Data were epoched in 2-second intervals and inspected using a semi-automated algorithm to remove epochs containing artefacts. Independent components analysis (ICA) was then used to remove eye blink and muscle artefacts. Following ICA, EEG data were re-referenced to the common average reference.

### Behavioural measures

Mood was assessed using the MADRS, a validated clinician-rated scale of depression symptom severity (Montgomery and Asberg, 1979). Response and remission were defined as a reduction in MADRS ≥ 50% from baseline, and a post-intervention MADRS total score < 10, respectively.

The visual 3-back task, adapted from Mull and Seyal (2001), was administered using Inquisit 4 software (Version 4, Millisecond Software) to assess working memory. Participants were asked to observe a sequence of letter stimuli presented on a computer screen in random order and respond (by pressing the spacebar on a keyboard) when the target letter currently on the screen matched the letter presented three trials previously (see Fig. 1C). The 3-back task is therefore similar to the Emotional Faces Memory Training task used for CET, but does not require processing of emotional content. The 3-back task consisted of 40 target letters interspersed with 180 non-targets, with a 2-second interval between successive letters. Prior to the start of each experiment, participants practiced the task for approximately 5 minutes to ensure they understood task instructions. The 3-back task was selected because it is considered challenging, requiring greater attentional and executive resources, and so reduces the likelihood of ceiling effects for participants that improved cognitively over the 6-week intervention. The working memory outcome measures were response time (RT) for correct responses, and d-prime, a measure of discriminate sensitivity (Haatveit *et al*., 2010).

### Neurophysiological measures

#### Resting-state power spectral density

Power spectra were calculated for resting-state EEG under eyes open and eyes closed conditions. Data from the first 15 seconds were discarded to ensure participants were completely at rest in the time window being analysed and power spectral densities (PSD) were estimated using 285 seconds of data. Log-normalised power spectral density values (μV^2^/Hz) were estimated for each EEG electrode over a range of 1 – 70Hz using the fast Fourier transform with 2-second sliding Hamming windows with 50% overlap, as initially described by Welch (1967).

Frontal alpha asymmetry was obtained under the eyes closed resting-state EEG condition (Van Der Vinne *et al*., 2017). To calculate this index, power in the alpha frequency band (8-13 Hz) was obtained at EEG channels F3 and F4. We then divided the difference in alpha power between channels by the sum of alpha power between channels: (F4 − F3)/(F4 + F3).

Frontal theta was calculated as the average of power in the theta frequency band (4-8 Hz) at anterior EEG channels Fp1, Fp2, Afz, F3, Fz, and F4 under the eyes closed resting-state EEG condition.

#### Event-related potentials

Event related potentials (ERPs) were calculated by averaging across target and non-target trials from the 3-back task. Post-stimulus activity was baseline-corrected using the mean amplitude from 500 ms to 0 ms prior to stimulus onset. Average amplitudes for ERP components P2 and P3 were measured from frontal EEG channel Fz. The time-window for averaging was determined by computing the grand average ERP component across BASELINE and POST periods for all participants and isolating the mean latency for the P2 and P3 components. The average amplitude was then calculated for each participant in a time-window ±20 ms around the grand average latencies for P2 and P3 components from the previous step. The latency of the P2 component was identified as 145.5 ms post-stimulus (time window for averaging: 125.5 – 165.5 ms), and the latency for the P3 component was 373.0 ms (time window for averaging: 353.0 – 393.0 ms) following stimulus onset.

#### Time-frequency analysis

Similar to ERP measures, time-frequency power was calculated by averaging across target and non-target trials from the 3-back task. We used a Hanning taper with a fixed 500 ms time window. Power values were baseline-corrected using activity from 500 ms to 0 ms prior to stimulus onset and transformed into a decibel scale (10*log10 of the signal). Alpha event-related desynchronization (ERD) was operationalised as band power 200 – 700 ms from stimulus onset in the alpha frequency band (8 – 12Hz). Theta event-related synchronisation (ERS) was defined as band power 0 – 500 ms from stimulus onset in the theta frequency band (4 – 8 Hz).

### Statistical analysis

Statistical analyses were performed using R statistical software version 3.5.1 (R Core Team, 2018) and the Fieldtrip MATLAB toolbox (Oostenveld *et al*., 2011). Scripts used for statistical analysis are available at the following link (https://github.com/snikolin/tDCSandCET). Two-tailed paired samples t-tests were performed for mood, working memory, and EEG neurophysiological outcomes to evaluate changes from BASELINE to POST. Normality of paired differences was tested using the Shapiro-Wilk test. A p-value of < 0.05 was considered non-normally distributed, in which case we additionally performed a non-parametric Wilcoxon-signed-rank test. We applied a Bonferroni correction to the six neurophysiological outcomes to reduce the false-positive (Type 1) error associated with multiple comparisons. As such, the threshold for statistical significance for EEG measures was set at *p* = 0.008 (i.e. 0.05/6).

Repeated measures and Spearman correlations between behavioural and neurophysiological measures are also provided in Supplementary Materials (see Supplementary Tables 1 and 2).

#### Equivalence tests

A within-subjects study design of n = 20 allows for the detection of moderate-large effect sizes of Cohen’s *d* ≥ 0.65 using α = 0.05 and 80% statistical power. Therefore, lack of statistical significance on a paired-samples t-test may be indicative of no effect or of insufficient power to detect effects sizes smaller than this threshold. To statistically reject the possibility of non-trivial effect sizes, we performed two one-sided t-test (TOST) equivalence procedures using Cohen’s *d* = 0.3 as the smallest effect size of interest. Thus, a significant equivalence test result would suggest an effect size of *d* < 0.3. This procedure was performed using the R package, TOSTER, and the *TOSTpaired* function (Lakens, 2017).

#### Non-parametric cluster-based permutation tests

In additional to the previous analyses for single electrodes (or averages of a small number of electrodes), we also conducted exploratory non-parametric cluster-based permutation analyses on the entire EEG dataset for measures extracted using power spectral density, event related potential, and time-frequency analyses. This method controls for multiple comparisons while assessing global changes from BASELINE to POST across all EEG channels, frequency ranges, and post-stimulus time points for event-related measures (i.e. ERPs and ERS/ERD); changes that were not otherwise hypothesised *a priori* (Maris & Oostenveld, 2007).

The Monte-Carlo method was used, whereby trials were randomly permuted in 3000 iterations and the resulting distributions were statistically compared using dependent samples *t*-tests. A two-tailed significance threshold of α < 0.05 was adopted as the cluster threshold and significance threshold for all analyses. Cluster-based permutation tests commonly require clusters of at least two neighbouring channels; however, due to the limited number of EEG channels in the current experiment, we opted for a less conservative threshold of at least one neighbouring channel.

For ERPs and ERS/ERD, the non-parametric cluster-based permutation tests were performed on a time interval of interest 0 – 1000 ms from the onset of 3-back task letter stimuli. For neurophysiological outcomes with frequency information, including resting-state PSDs and ERS/ERD, permutation testing was conducted in the frequency band of 1 – 70Hz.

## RESULTS

A total of twenty participants with treatment resistant depression completed 18 sessions of tDCS combined with CET. Table 1 shows baseline demographic and clinical information for the sample.

**Table 1.**
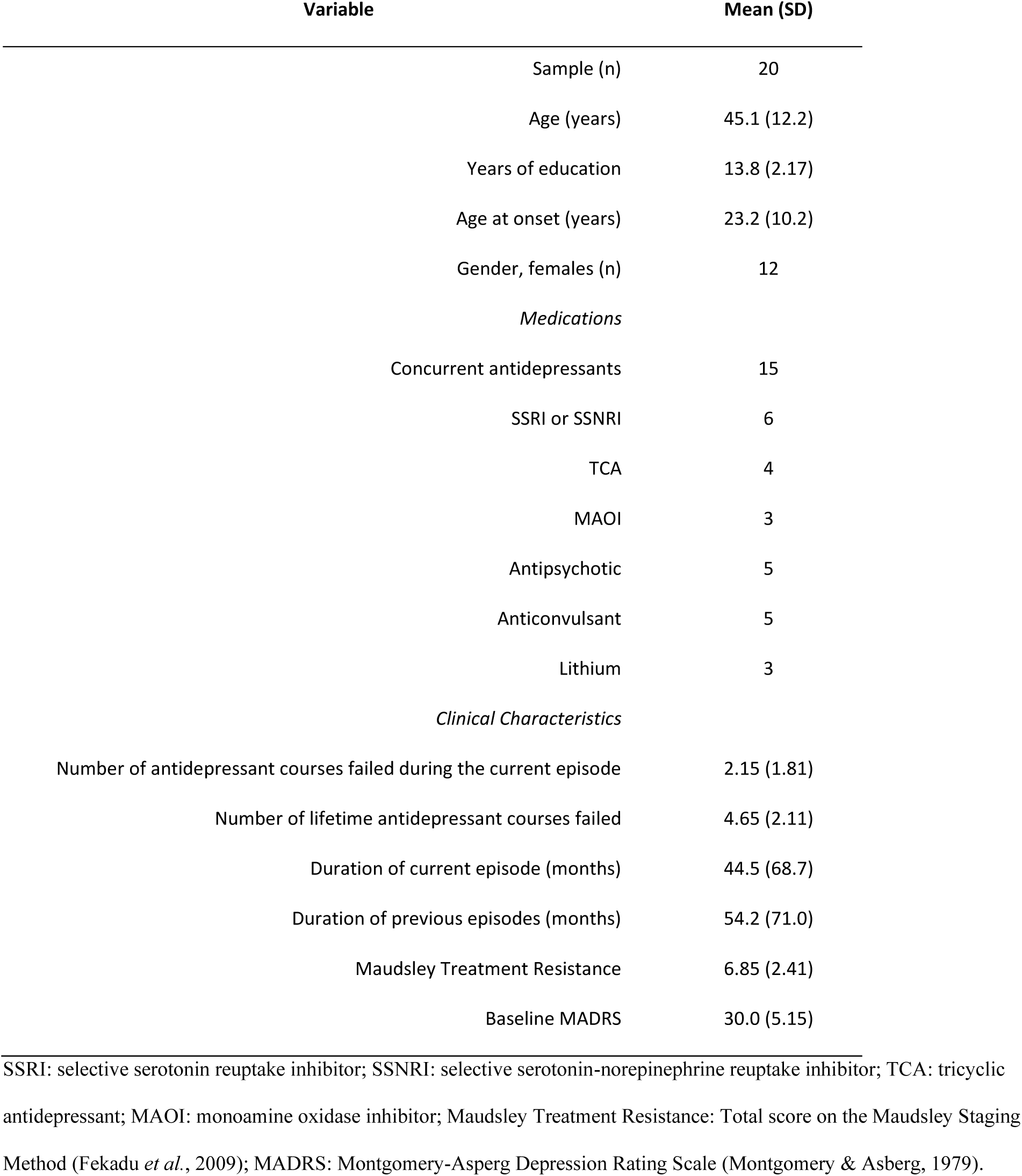
Baseline demographic and clinical information.

| Variable | Mean (SD) |
| --- | --- |
| Sample (n) | 20 |
| Age (years) | 45.1 (12.2) |
| Years of education | 13.8 (2.17) |
| Age at onset (years) | 23.2 (10.2) |
| Gender, females (n) | 12 |
| *Medications* |  |
| Concurrent antidepressants | 15 |
| SSRI or SSNRI | 6 |
| TCA | 4 |
| MAOI | 3 |
| Antipsychotic | 5 |
| Anticonvulsant | 5 |
| Lithium | 3 |
| *Clinical Characteristics* |  |
| Number of antidepressant courses failed during the current episode | 2.15 (1.81) |
| Number of lifetime antidepressant courses failed | 4.65 (2.11) |
| Duration of current episode (months) | 44.5 (68.7) |
| Duration of previous episodes (months) | 54.2 (71.0) |
| Maudsley Treatment Resistance | 6.85 (2.41) |
| Baseline MADRS | 30.0 (5.15) |

SSRI: selective serotonin reuptake inhibitor; SSNRI: selective serotonin-norepinephrine reuptake inhibitor; TCA: tricyclic antidepressant; MAOI: monoamine oxidase inhibitor; Maudsley Treatment Resistance: Total score on the Maudsley Staging Method (Fekadu *et al.*, 2009); MADRS: Montgomery-Asperg Depression Rating Scale (Montgomery & Asberg, 1979).

### Behavioural outcomes

A summary of statistical outcomes for behavioural measures is provided in Table 2. tDCS combined with CET was found to significantly reduce symptoms of depression (*p* < 0.001). Eight participants met the response criterion (40%), and three participants were remitters upon the completion of 18 sessions of treatment (15%). Mood improvements persisted at the 1-month follow-up (n = 14, t_13_ = -3.56, p = 0.003, d = 0.94), but were no longer significant at the 3-month follow-up, in which only half of the original sample remained (n = 10, t_9_ = -2.17, p = 0.06, d = 0.69). Participants significantly improved on working memory d-prime (*p* = 0.03), but not response time (*p* = 0.45; see Fig. 2).

**Table 2.** Summary of statistical outcomes for behavioural and neurophysiological measures. Baseline and post-intervention summary measures are shown for mood, working memory, and EEG outcomes. Paired samples *t*-tests were used to examine changes from baseline to post. Equivalence tests were used to determine whether effects were significantly lower than the smallest effect size of interest i.e. an effect size of Cohen’s *d* = 0.3.

| Test |  | Baseline |  | Post |  | Post – Baseline |  | Paired Samples T-Test |  |  | Equivalence Test |  |
| --- | --- | --- | --- | --- | --- | --- | --- | --- | --- | --- | --- | --- |
|  |  | Mean (SD) |  | Mean (SD) |  | Mean (95% CI) |  | t<sub>19</sub> | p | Cohen's d | t<sub>19</sub> | p |
| Mood Outcome |  |  |  |  |  |  |  |  |  |  |  |  |
|  | MADRS | 30.0 | 5.0 | 19.5 | 9.5 | -10.5 | -5.7 – -15.2 | -4.61 | <0.001 | 1.03 | 3.27 | 0.998 |
| Working Memory |  |  |  |  |  |  |  |  |  |  |  |  |
|  | D-prime | 1.62 | 0.76 | 2.01 | 0.84 | 0.39 | 0.05 – 0.73 | 2.40 | 0.027 | 0.54 | -1.06 | 0.848 |
|  | RT (ms) | 796 | 174 | 835 | 235 | 40 | -68 – 148 | 0.77 | 0.452 | 0.17 | 0.57 | 0.287 |
| Power Spectral Density |  |  |  |  |  |  |  |  |  |  |  |  |
|  | Frontal Alpha Asymmetry | 0.09 | 0.31 | 0.04 | 0.25 | -0.05 | -0.24 – 0.14 | -0.55 | 0.591 | 0.12 | -0.80 | 0.218 |
|  | Frontal Theta | -0.18 | 1.00 | -0.09 | 1.17 | 0.09 | -0.44 – 0.62 | 0.35 | 0.729 | 0.08 | 0.99 | 0.167 |
| Event-Related Potentials |  |  |  |  |  |  |  |  |  |  |  |  |
|  | P2 | 1.64 | 1.27 | 2.06 | 1.46 | 0.42 | -0.004 – 0.85 | 2.07 | 0.052 | 0.46 | -0.73 | 0.763 |
|  | P3 | 1.01 | 1.65 | 1.04 | 1.27 | 0.03 | -0.41 – 0.47 | 0.31 | 0.888 | 0.03 | 1.20 | 0.123 |
| Time-Frequency Representations |  |  |  |  |  |  |  |  |  |  |  |  |
|  | Alpha ERD | -1.24 | 1.43 | -1.35 | 1.31 | -0.11 | -0.51 – 0.29 | -0.57 | 0.575 | 0.13 | -0.77 | 0.225 |
|  | Theta ERS | 0.46 | 0.69 | 0.51 | 0.83 | 0.05 | -0.18 – 0.29 | 0.47 | 0.641 | 0.11 | 0.87 | 0.198 |

MADRS: Montgomery-Asperg Depression Rating Scale (Montgomery & Asberg, 1979); ERD: Event-related desynchronization; ERS: Event-related synchronisation.

**Figure 2.**
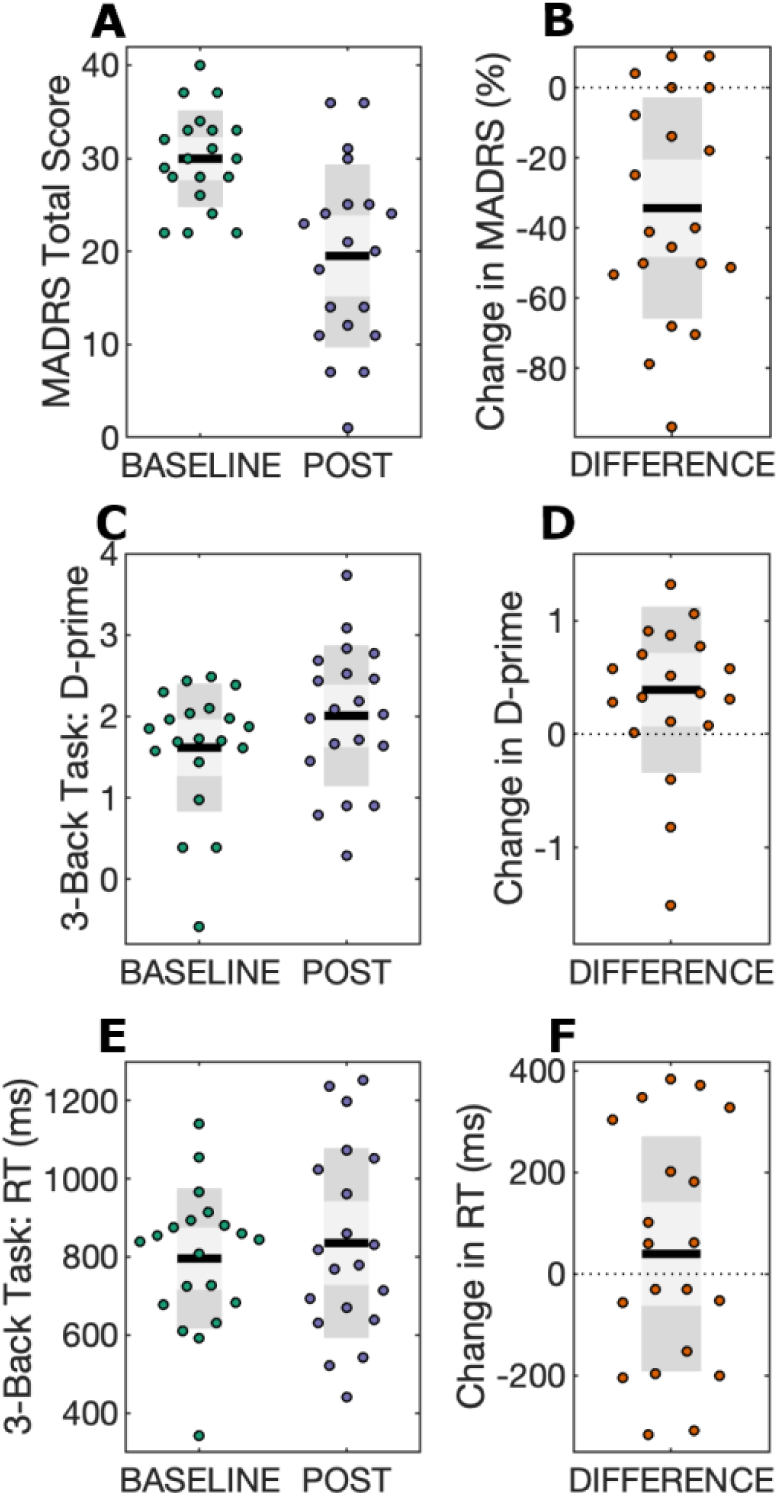
Behavioural outcomes. Scatter plots for mood and working memory performance scores are displayed at BASELINE and POST time points, in addition to change scores (DIFFERENCE). Black lines show the mean, light grey shaded boxes indicate the standard deviation, and dark grey regions indicate the 95% confidence interval. **A)** Total MADRS scores; **B)** MADRS difference scores, calculated as the percentage change in MADRS using the equation (POST – BASELINE)/BASELINE; **C)** 3-back task d-prime scores; **D)** D-prime difference scores, calculated as POST – BASELINE; **E)** 3-back task response time (RT) scores; **F)** RT difference scores, calculated as POST – BASELINE.

### Neurophysiological outcomes

A summary of statistical outcomes for neurophysiological measures is provided in Table 2. Frontal alpha asymmetry, frontal theta, and theta ERS were not normally distributed, thus additional Wilcoxon-signed-rank tests were performed.

#### Resting-state power spectral density

There were no significant effects for frontal alpha asymmetry (*p* = 0.59; Wilcoxon-signed-rank test, *p* = 0.70) or frontal theta (*p* = 0.73; Wilcoxon-signed-rank test, *p* = 0.96) change from BASELINE to POST time periods. Equivalence testing for both measures was not significant, thus an effect size greater than *d* = 0.3 cannot be statistically rejected. Exploratory cluster-based permutation testing did not reveal any significant spectral power changes for any frequency band for eyes-closed or eyes-open resting-state EEG conditions (see Fig. 3).

**Figure 3.**
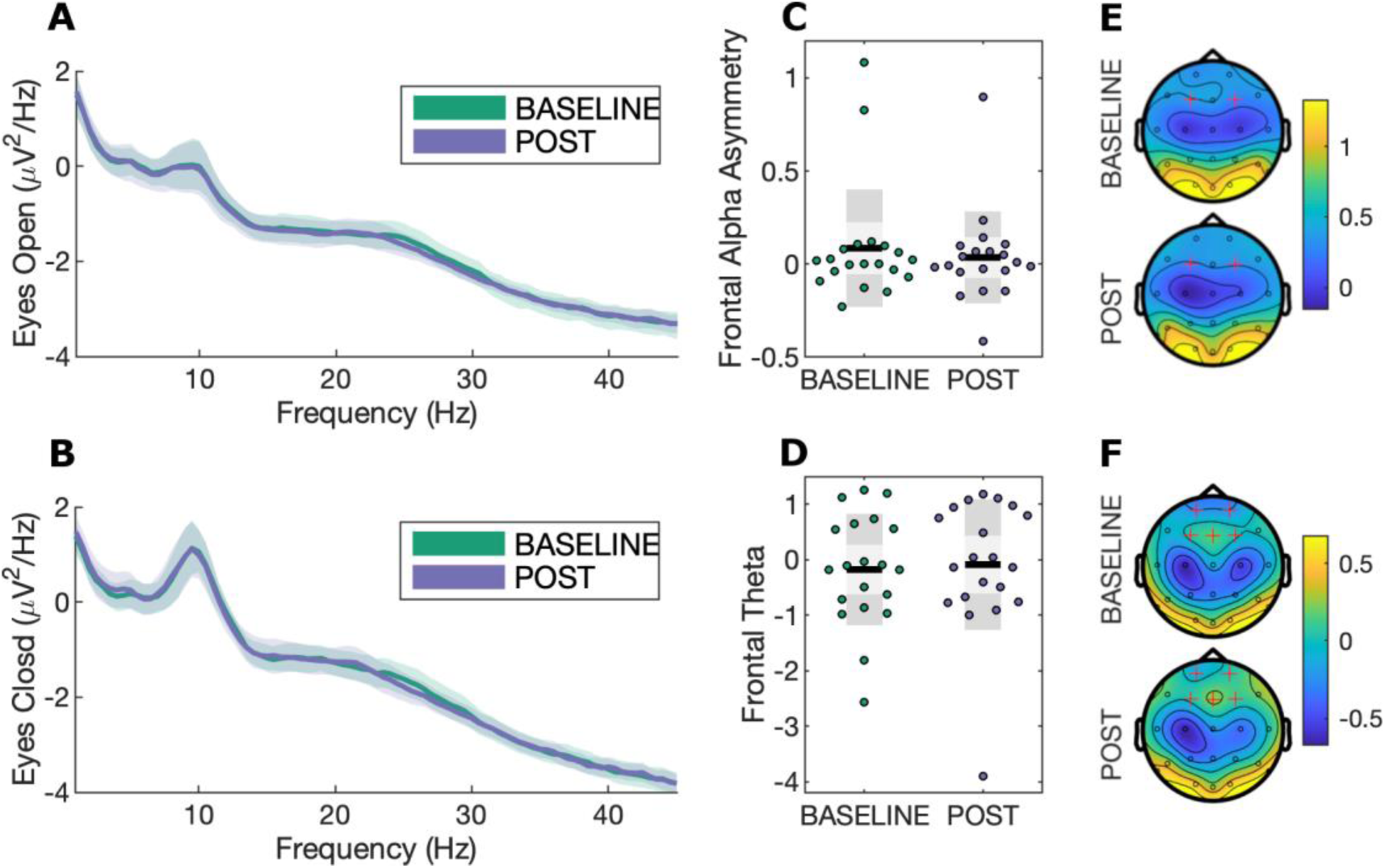
Resting-state power spectral density. **A)** Power spectra for eyes open resting-state EEG with bootstrapped 95% confidence intervals at channel Cz; **B)** Power spectra for eyes closed resting-state EEG with bootstrapped 95% confidence intervals at channel Cz; **C)** Frontal alpha asymmetry scatterplot during eyes-closed resting-state EEG, calculated as [F4 − F3]/[F4 + F3]. Black lines show the mean, light grey shaded boxes indicate the standard deviation, and dark grey regions indicate the 95% confidence interval; **D)** Frontal theta during eyes-closed resting state; **E)** Topography for alpha (8-13Hz) power. Highlighted channels (F3 and F4) were used to calculate the frontal alpha asymmetry measure; **F)** Topography for theta (4-8Hz) power. Highlighted channels were averaged to calculate the frontal theta index.

#### Event-related potentials

There was no significant effect for the P3 ERP component (*p* = 0.89). There was a moderate increase in P2 amplitude (*p* = 0.05), however, this did not survive correction for multiple comparisons using the Bonferroni adjusted *p*-value threshold (i.e. *p* < 0.008). Equivalence testing for P2 and P3 components was not significant. No significant clusters were observed comparing ERP measures at BASELINE to POST (see Fig. 4).

**Figure 4.**
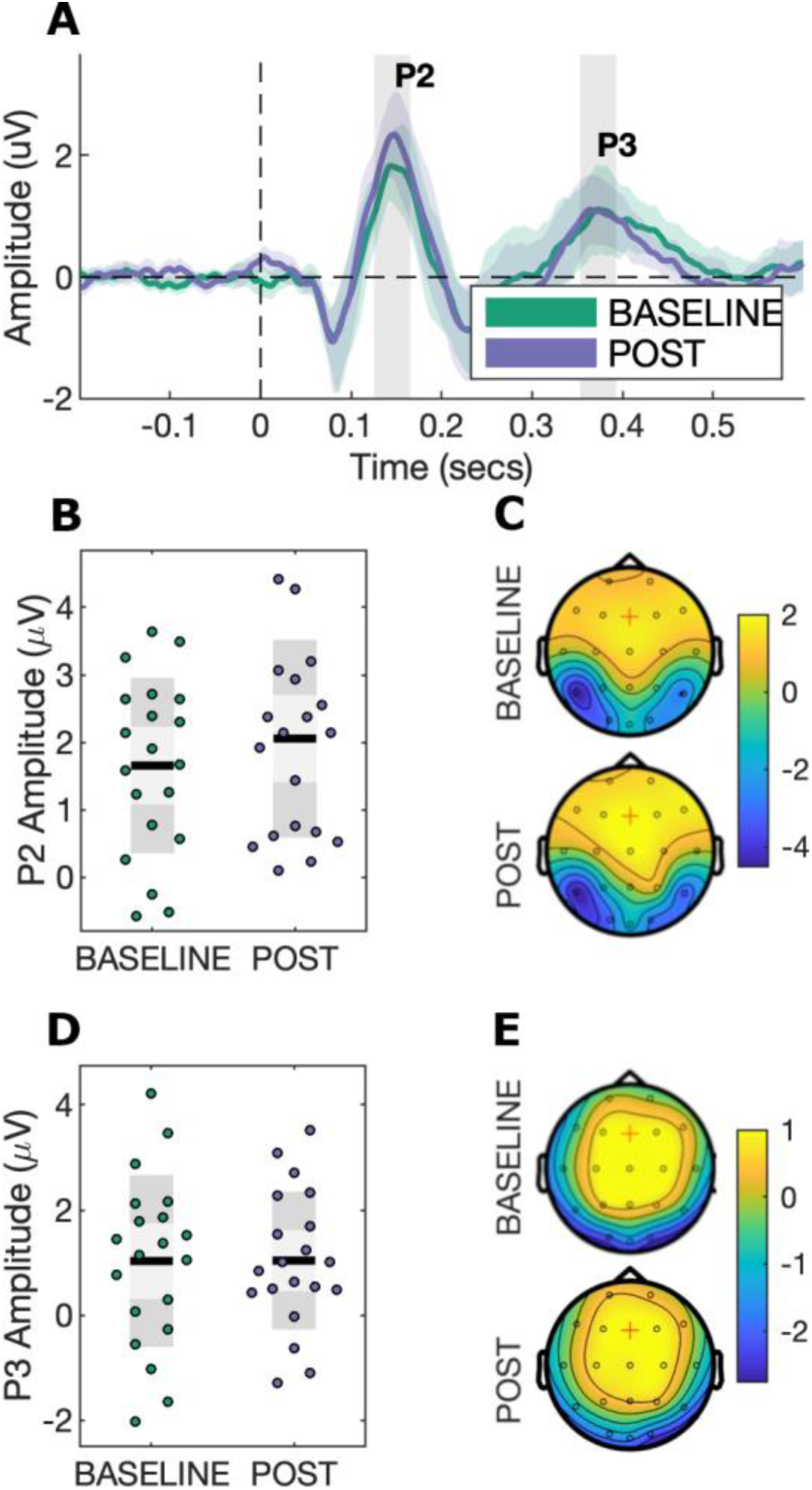
Event-related potentials during the 3-back task. **A)** ERPs with bootstrapped 95% confidence intervals at channel Fz; **B)** P2 component scatterplot. Black lines show the mean, light grey shaded boxes indicate the standard deviation, and dark grey regions indicate the 95% confidence interval; **C)** Topography for the P2 component (145.5 ms ± 20 ms); **D)** P3 component scatterplot; **E)** Topography for the P3 component (373 ms ± 20 ms).

#### Time-frequency analysis

There were no significant effects for alpha ERD (*p* = 0.58) or theta ERS (*p* = 0.64; Wilcoxon-signed-rank test, *p* = 0.93). Likewise, equivalence testing was not significant for both measures. No significant clusters were observed at any frequency bands, channels, or time points (see Fig. 5).

**Figure 5.**
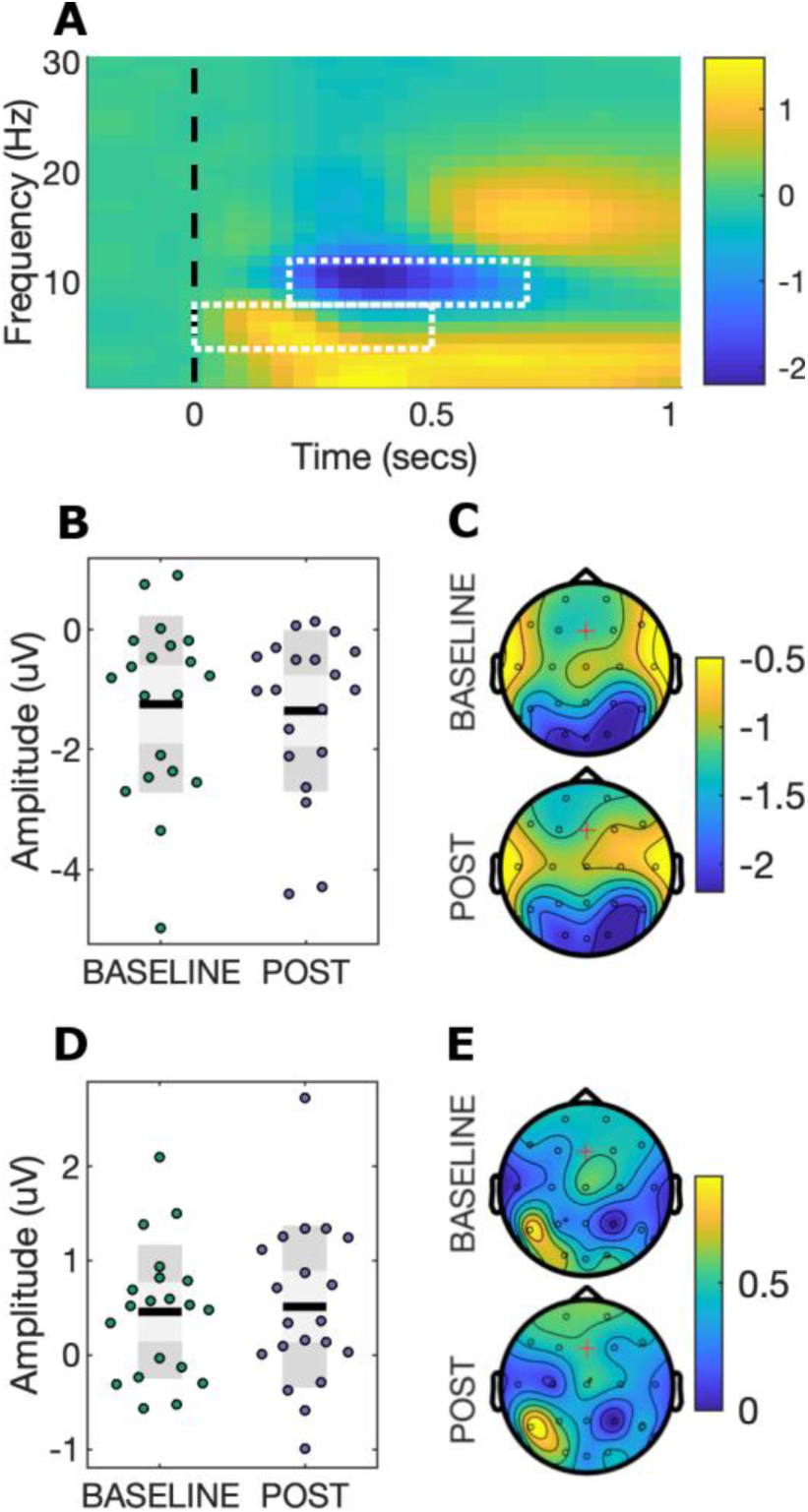
Time-frequency power during the 3-back task. **A)** EEG power at channel Fz. White boxes indicate regions of interest for alpha ERD and theta ERS; **B)** Alpha ERD scatterplot. Black lines show the mean, light grey shaded boxes indicate the standard deviation, and dark grey regions indicate the 95% confidence interval; **C)** Topography for alpha ERD; **D)** Theta ERS scatterplot; **E)** Topography for theta ERS.

### Safety

Safety was assessed using a side effects questionnaire adapted from Brunoni *et al*. (2011), which has been used in our prior studies (e.g. Loo *et al*. 2011). The combined tDCS and CET intervention was well-tolerated by participants and resulted in transient adverse events of mild-moderate severity. The most frequent side effects were skin redness, paraesthesia (tingling, burning, and itching), and headache, in agreement with prior meta-analyses of tDCS adverse events (Moffa *et al*., 2017; Nikolin *et al*., 2018a). As reported in our previous work examining the cognitive effects of tDCS with concurrent CET, treatment did not result in significant reductions in cognitive functioning (Martin *et al*., 2018). A summary of all adverse events is provided in Table 3.

**Table 3.** Adverse events. The group-level estimate of adverse event frequency is provided using the mean, standard deviation (SD), and range of sessions in which an adverse event occurred. Incidence rates are also reported as a percentage (%) of sessions in which an adverse event occurred out of the total of all tDCS and CET sessions conducted for the study.

| Adverse event | Frequency (#) |  | Incidence (%) |
| --- | --- | --- | --- |
|  | Mean | (SD; range) |  |
| Redness | 10.7 | (6.2; 0 – 18) | 59.4 |
| Tingling | 6.7 | [6.5; 0 – 18) | 37.2 |
| Burning | 4.3 | (5.5; 0 – 18) | 23.9 |
| Itching | 2.8 | (3.8; 0 – 16) | 15.6 |
| Headache | 1.9 | (2.6; 0 – 7) | 10.3 |
| Fatigue | 1.5 | (2.1; 0 – 8) | 8.1 |
| Others | 1.2 | (2.2; 0 – 8) | 6.4 |
| Blurred vision | 0.9 | (3.6; 0 – 16) | 4.7 |
| Light-headedness | 0.4 | (0.7; 0 – 2) | 2.2 |
| Nausea | 0.4 | (1.3; 0 – 6) | 1.9 |
| Pain | 0.3 | (0.7; 0 – 2) | 1.7 |

## DISCUSSION

This study investigated the neurophysiological effects of tDCS combined with CET in participants with treatment resistant depression. Following the intervention, participants improved in mood as well as working memory performance accuracy, but not response time, on a visual 3-back task. Contrary to our hypotheses, for EEG outcomes, there was no evidence of differences in resting-state measures (frontal alpha asymmetry and frontal theta), or in task-related measures (alpha ERD, theta ERS, and the P3 ERP component). We found weak evidence to suggest that P2 amplitude, a marker of attentional processes, increased, but this result did not survive correction for multiple comparisons.

Combining tDCS with CET appears to be a promising augmentation strategy to improve therapeutic efficacy in treatment resistant depression. In a clinical pilot study we found that this combined intervention was feasible, safe, and associated with significant antidepressant effect (Martin *et al*., 2018). Accompanying the improvement in mood, we observed a near transfer effect from CET to improved discriminate sensitivity (i.e. d-prime) on the 3-back working memory task in the current report. In a similar study combining five repeated sessions of tDCS with cognitive control training, Segrave *et al*. (2014) also noted improved accuracy on an affective 2-back task. Our findings are in line with near transfer effects following working memory training for improving cognition (Minear *et al*., 2016; Soveri *et al*., 2017). A meta-analysis by Schwaighofer *et al*. (2015) suggests that the near transfer effect of working memory training has an effect size ranging between Hedge’s *g* = 0.37-0.72, which encompasses the effect size for d-prime observed in the present study (Cohen’s *d* = 0.54). In the absence of a control condition, however, it is difficult to determine whether working memory improvements observed in the current study are due to the intervention (i.e. near transfer effects from CET), or may be associated with mood improvement.

Although there were no significant changes in neurophysiological measures after controlling for multiple comparisons, we found weak evidence that the intervention increased the P2 ERP amplitude. The P2 component has been linked to attentional processes (Kemp *et al*., 2006; Lijffijt *et al*., 2009; Luu *et al*., 2014; Yuan *et al*., 2016; Vilà-Balló *et al*., 2018)), whereas P3 is associated with higher-order cognitive functions such as target identification and categorisation (Friedman *et al*., 2001; Kok, 2001; Rac-Lubashevsky & Kessler, 2016). McEvoy *et al*. (1998) systematically investigated the impact of working memory load and practice effects, and similarly observed changes in P2, but not P3, amplitude. Our results therefore suggest that CET, which involves a working memory component, may improve lower order processing and attentional capacity, observed as an increase in P2 amplitude, although the statistical evidence was only weak. In line with this interpretation, similar findings were obtained following 3 weeks of treatment for depression using transcranial magnetic stimulation, another form of non-invasive brain stimulation, which increased P2 ERP amplitudes during an oddball task (Choi *et al*., 2014). Likewise, Spronk *et al*. (2008) observed a left lateralised increase in frontal P2 amplitude in depressed patients during an auditory oddball task following 20 treatment sessions of transcranial magnetic stimulation, providing additional support for improvements in sensory-attentional processes but not higher-order cognitive functions. Our results are also in agreement with Keeser et al., (2011), who investigated the acute after-effects of prefrontal tDCS on working memory functioning. They found an increase in P2 amplitude at midline frontal electrode Fz, in conjunction with improved d-prime accuracy and reduced reaction times. However, it is important to note that these findings may reflect differing neuromodulatory processes compared to the long-term, cumulative effects investigated in the current study.

Resting-state EEG measures commonly associated with depression, such as frontal alpha asymmetry and frontal theta power, were not significantly altered by the treatment despite meaningful improvements in mood. These measures were hypothesised *a priori* to change with treatment because they had been linked to antidepressant response (Spronk *et al*., 2011; Arns *et al*., 2015; Arns *et al*., 2016). Recently, there have been some criticisms of the viability of alpha asymmetry as a relevant factor for depression (Van Der Vinne *et al*., 2017). Debener *et al*. (2000) challenge the notion that alpha asymmetry is a trait marker for depression, and report that increased variability, rather than increased alpha lateralisation, may be a more relevant measure. Similarly, frontal theta PSD may be better interpreted as a trait measure, reflecting structural or function abnormalities that contribute to the risk of developing depression, rather than a biomarker of the state of being depressed. Indeed, Hunter *et al*. (2013) found theta generated in the rostral anterior cingulate cortex to be a trait predictor of depression that does not change with treatment i.e. it is invariant with regards to an individual’s diagnosis of depression. Interestingly, supplementary analyses show a correlation between frontal alpha asymmetry (*p*=0.029) and mood scores (see Supplementary Table 1). However, these must be interpreted with caution as correlations using small samples have a low probability of replication and can result in exaggerated effects through chance alone. Monte-Carlo simulations show that correlations converge, that is, produce stable estimates close to the true effect, only as sample sizes approach n=250 (Schönbrodt & Perugini, 2013).

Our lack of positive neurophysiological findings is generally consistent with the tDCS literature for protocols using repeated sessions of stimulation. To the best of our knowledge, only three studies have identified functional or structural changes associated with repeated sessions of anodal tDCS to the left DLPFC. Previous work by our group found an increase in neuroplasticity, quantified by probing motor cortex reactivity using transcranial magnetic stimulation, following prefrontal tDCS treatment for depression (Player *et al*., 2014). In another clinical application, Ulam *et al*. (2015) examined the neurorehabilitatory effects of 10 consecutive sessions of left DLPFC tDCS for traumatic brain injury. They did not observe any significant effects from baseline to the final session of stimulation on EEG measures, including an analysis of frontal (F3) theta power, but did note a significant interaction effect compared to sham tDCS for delta and alpha frequency bands. Mondino *et al*. (2015) delivered 10 sessions of anodal stimulation over the left DLPFC, with cathodal stimulation over the temporo-parietal junction, for the treatment of auditory verbal hallucinations in schizophrenia and showed increased resting-state functional connectivity in the active tDCS group compared to sham. The present study did not include a sham control group, and thus we were unable to examine interaction effects, which may have revealed similar neurophysiological changes.

There may be several reasons why we did not observe significant neurophysiological changes in the present study. Firstly, the modest sample size only allowed for detection of moderate-large effects. The estimated effect sizes for the EEG measures were small (*d* = 0.03 − 0.13), with the exception of P2 amplitude (*d* = 0.46). Although the majority of these effects were less than the smallest effect size of interest (*d* = 0.3), equivalence testing was non-significant for all neurophysiological outcomes. This suggests that there was sufficient variability in participant outcomes that the true effect could not be identified as less than our threshold criteria for equivalence testing. Thus, we are also unable to rule out the possibility of effect sizes greater than Cohen’s *d* = 0.3. Larger sample sizes are therefore required to overcome issues of heterogeneity, and associated variability, inherent in depressed samples, and to thereby improve the ability to draw conclusions from statistical testing. For example, our results indicate that a sample of 39 participants is required to detect the effect size for P2 amplitude (*d* = 0.46) with 80% power.

Another potential reason for our null findings could be that the neurophysiological measures selected for hypothesis testing were suboptimal to detect treatment effects. To counteract this limitation, we examined resting-state PSD, as well as ERP and ERS/ERD during the 3-back task, using exploratory cluster-based non-parametric permutation tests to account for the possibility that an effect may lie beyond our *a priori* hypotheses, and found no significant clusters. Nevertheless, more sophisticated analyses, such as theta-gamma phase-amplitude coupling (Sun *et al*., 2015; Noda *et al*., 2017), or dynamic causal modelling to determine changes in effective connectivity between brain regions (Lu *et al*., 2012; Harding *et al*., 2015; Breakspear, 2017), may reveal the effects of the intervention. For example, theta connectivity localised to the anterior cingulate cortex has been identified as a predictor of antidepressant response to treatment courses using transcranial magnetic stimulation (Narushima *et al*., 2010; Bailey *et al*., 2018), and sertraline (Pizzagalli *et al*., 2018).

Lastly, depression has been linked to abnormal activity in the subgenual anterior cingulate cortex, an area of the limbic system associated with emotional information processing (Pizzagalli *et al*., 2002; Drevets *et al*., 2008; Matthews *et al*., 2008; Rentzsch *et al*., 2014). The Emotional Faces Memory Training task used for CET in the current study was designed to concurrently activate the cognitive control network and increase brain activity in these deeper, limbic regions. Therefore, the effects of the intervention may be more readily observed within these subcortical structures. However, identification of changes at this depth may lie beyond the reach of the EEG setup used in this experiment. Though other EEG studies have successfully localised dysfunctional brain activity to sources within the anterior cingulate (Wacker *et al*., 2009), this is typically achieved using EEG systems with a larger number of recording channels to better estimate source activity. It is also possible that greater effects would have been observed using task-related electrophysiological measures during an emotional processing task similar to the CET, such as passive viewing of emotionally salient images (MacNamara *et al*., 2016). Stewart *et al*. (2014) found that alpha asymmetry was a better predictor when measured during an emotional task rather than during resting-state EEG. Unfortunately, this data was not collected, but may be a valuable addition to future studies seeking to examine similar interventions in depression.

## CONCLUSION

Combining tDCS with CET is a promising augmentation strategy for treatment resistant depression. Although behavioural measures identified improvements in mood and working memory accuracy, the neurophysiological mechanisms underlying these improvements remain elusive. EEG analyses of resting-state (i.e. PSD) and task-related activity (i.e. ERPs and ERS/ERD) did not reveal substantial changes from baseline to post-treatment. There is, however, tentative evidence of an increase in P2 amplitude, suggesting that tDCS and CET may improve attention modulation and context updating in depressed participants. These findings do not rule out the possibility that more sophisticated EEG analyses, such as phase-amplitude coupling or functional/effective connectivity, may uncover significant effects and better explain observed antidepressant and working memory changes. Further research is needed using a sham control condition and a larger sample of participants to confirm our preliminary neurophysiological results and better quantify the mechanisms of action for this promising new intervention.

## ACKNOWLEDGEMENTS

Donel Martin and Tjeerd Boonstra were both funded by a NARSAD Young Investigator Award (grant numbers 24015 and 26060, respectively) from the Brain and Behavior Research Foundation.

## AUTHOR CONTRIBUTIONS

Designed the Study: DM, TB, CL; Performed Experiments: SN; Analyzed Data: SN, DM, TB; Interpreted Results: SN, DM, TB, CL, BI; Prepared Figures: SN; Drafted Manuscript: SN; Edited/Revised Manuscript: SN, DM, CL, BI, TB; Approved the Final Manuscript: SN, DM, CL, BI, TB.

## DATA ACCESSIBILITY STATEMENT

Data, MATLAB and R scripts used for EEG processing, calculation of neurophysiological measures, and statistical analyses are available at the following link: https://github.com/snikolin/tDCSandCET.

## ABBREVIATIONS

CET: cognitive emotional training
DLPFC: dorsolateral prefrontal cortex
EEG: elecetroencephalography
EFMT: emotional faces memory task
ERD: event related desynchronization
ERP: event related potential
ERS: event related synchronisation
ICA: independent component analysis
MADRS: Montgomery-Asperg depression rating scale
MDD: major depressive disorder
PSD: power spectral density
RT: response time
tDCS: transcranial direct current stimulation
TOST: two one-sided *t*-test

